# Tomographic-encoded multiphoton (TEMP) microscopy

**DOI:** 10.1101/2022.04.11.487875

**Authors:** Hongsen He, Xin Dong, Yu-Xuan Ren, Cora S. W. Lai, Kevin K. Tsia, Kenneth K. Y. Wong

**Affiliations:** Department of Electrical and Electronic Engineering, The University of Hong Kong, Pokfulam Road, Hong Kong, China; School of Biomedical Sciences, Li Ka Shing Faculty of Medicine, The University of Hong Kong, Pokfulam Road, Hong Kong, China; Institute for Translational Brain Research, Shanghai Medical College, Fudan University, Shanghai, China; State Key Laboratory of Brain and Cognitive Sciences, The University of Hong Kong, Pokfulam Road, Hong Kong, China; Advanced Biomedical Instrumentation Centre, Hong Kong Science Park, Shatin, New Territories, Hong Kong, China

## Abstract

Axial scanning in multiphoton microscopy (MPM) is typically realized by mechanically shifting either the objective or the sample. However, the scan speed is usually hindered by the mechanical inertia of the bulky mass. Although the extended depth of field provided by the non-diffracting beam allows fast volumetric imaging, it abandons the axial resolution. Here, we demonstrate a novel and powerful tomographic technique using the Bessel droplet in MPM, termed Tomographic-Encoded MultiPhoton (TEMP) microscopy. We show that benefiting from the high-order nonlinear excitation in MPM, the side-lobes cancellation and smaller beam focus of the Bessel droplet realize better image quality. The TEMP microscopy allows fast axial scanning, less risks of photodamage and photobleaching, and high-resolution and high-contrast imaging. Furthermore, fewer raw images are required for the 3D image reconstruction. To demonstrate its usability and advantages for scattering tissues and biomedical applications, we showcase the TEMP microscopy with highly scattering fluorescence microspheres and mouse brain slice. More details can be visualized by the Bessel droplet compared with the conventional Gaussian and Bessel beam. More importantly, the TEMP technique is an easy-plug-in method for the current microscopy system. The TEMP microscopy is promising for fast volumetric multiphoton imaging, especially for highly scattering tissues.

## Introduction

Multiphoton microscopy (MPM) stands out among optical microscopy technologies, mainly attributed to its unique ability to provide high spatial resolution and contrast at an unprecedented scale and depth inside the scattering tissues [1-5]. This attribute makes it ideal for a range of life science studies, such as in vivo animal imaging [6] and neuroscience [3, 7]. As excitation in MPM occurs only at the focal point of a diffraction-limited microscopy [8], the volumetric imaging of a thick biological specimen is typically realized by scanning the focus in three dimensions (3D). Individual optical sections are classically acquired in a raster-scan manner in the *x*-*y* plane, utilizing a pair of galvanometric mirrors, which can achieve a 2D frame rate up to 30 Hz [9], while a full 3D image is composed by serially scanning the specimen at sequential *z* positions. In most MPM modalities, the volumetric imaging speed is limited by the scanning rate of the *z*-axis. Approaches to achieve the axial scan can be typically classified into two groups. The first one is to mechanically shift the objective or the sample using a piezoelectric scanner [10] or motorized actuator [11]. However, the volume rate is typically limited to 1–10 Hz due to mechanical inertia [1]. The other approach is to move the laser focus axially without inertia. For example, using an electrically tunable lens (ETL) [12] or tunable acoustic gradient-index (TAG) [13, 14] lens to control focal shifts, but they discard higher-order aberrations, hindering to achieve a diffraction-limited focus. More recently, a series of *z*-scan techniques have been reported, including the reverberation loop for multiplexing laser pulses [15], light beads microscopy [16], hybrid multiplexed sculpted light microscopy [17], multi-Z confocal microscopy [18], and axial focusing converted by lateral scanning [19]. Notably, major modifications of current microscopies are required to apply these techniques. As such, there remains a need for an easy-plug-in technology to achieve fast axial scan in current microscopies.

Another inertia-free strategy is to exploit a non-diffracting beam (NDB, e.g., Bessel [20, 21] or Airy beam [22, 23]) with an extended depth of field (EDOF), capturing a 2D projected view of a volumetric structure [24-26]. This approach not only provides a fast speed volumetrically [20], but also has less attenuations in tissues owing to the self-healing nature of the NDB [22, 27]. However, its major limitation is the lack of axial resolution. Recently, a variety of methods have been adopted to resolve the depth based on the NDB, such as the mirrored Airy beams [23], V-shaped point spread function (PSF) [28], or rotating the sample [29]. The trade-offs of the first two approaches are the reduced field of view (FOV) and the requirements of sparse samples because of the co-existed two images. The third method has a complex optical alignment, and some samples, like liquid solutions, are not suitable to be rotated. In 2021, Gong *et al*. reported a depth-resolving method based on an optical beating technique (OBT) in stimulated Raman scattering (SRS) microscopy, where the volumetric information can be decoded from the spatial frequency domain, providing at least twofold improvement in imaging depth [27]. The beam used in OBT is also known as the Bessel droplet [30, 31], which maintains the nature of the NDB. The axial scanning speed is mainly determined by the refresh rate of the spatial light modulator (SLM). A state-of-the-art SLM now can achieve a speed up to 1 kHz, and if considering the faster digital micromirror device (DMD), the speed can be boosted to ∼20 kHz. One more advantage is that this approach does not need major adjustments to the current microscopy system, just by inserting the SLM or DMD as a reflecting mirror.

The OBT holds great potential for MPM, except for the high axial scanning speed. First, for conventional Gaussian beam scanning, the way to improve the signal-to-noise ratio (SBR) of the image is to increase the excitation intensity. However, as the fluorophore response is nonlinear, above the saturation threshold, little extra fluorescence signal is gained by rising illumination levels [32]. More importantly, high illumination power raises the risks of photodamage and photobleaching on fluorescence samples, especially for in vivo multiphoton imaging. In contrast, multiple illuminations using Bessel droplets increase the exposure time for each axial position while maintaining low illumination levels. Thus, it enhances the image SBR while lowers the above risks [33, 34]. Second, the side lobes of the Bessel droplet are serious, especially in the gap region [31], which greatly degrades the image quality and 3D reconstruction accuracy. Multiphoton excitations will eliminate the influence of the side-lobes owing to the nonlinearity-induced stronger excitation localization and reduced out-of-focus background.

In this work, we extend OBT to MPM, termed Tomographic-Encoded MultiPhoton (TEMP) microscopy. We show that OBT not only benefits from the side-lobes cancellation and smaller beam focus of the droplet for better image quality, but that the high-order nonlinear excitation leads to a qualitatively different performance. First, we show that, unlike one-photon (1P) excitation, the TEMP microscopy has better purity of the Bessel droplet, allowing more accurate reconstruction of the 3D image. Secondly, multiple illuminations from a series of Bessel droplets enhance the image SBR with a better resolution, leading to a superior visualization of weak fluorescence neurons. Thirdly, the image reconstruction of the TEMP microscopy requires fewer scans than the SRS OBT; furthermore, compared with the two-beam-overlapped SRS OBT, TEMP microscopy only needs a route of Bessel droplet. This means the axial scanning speed can be further improved in TEMP microscopy and the beam alignment is easier. We demonstrate the applicability of TEMP microscopy with highly scattering fluorescence microspheres and mouse brain slice.

### Working principle of TEMP microscopy

A Bessel beam can be regarded as a superposition of plane waves propagating with angle *θ* along the propagation direction (*z*-axis). The electric field of the 0th-order Bessel beam can be written in cylindrical coordinates as [35]

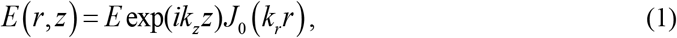

where *E* is the amplitude, *J*_0_ is the Bessel function of the first kind of order zero, while *k*_z_ and *k*_*r*_ are the wave vectors along the *z*- and *r*-axis, respectively. The Bessel droplet is generated by axially interfering two Bessel beams [Fig. 1(a)]. The electric field of two axially superposed Bessel beams is expressed as

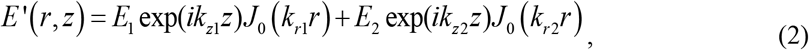

where suffixes 1 and 2 represent respective Bessel beams. Here, no relative phase shift between the two beams. When *E*_1_ is equal to *E*_2_, the axial intensity gradient of the Bessel droplet is the maximum [36], which will benefit the frequency modulation accuracy in the TEMP microscopy. The intensity distribution of the Bessel droplet along *z*-axis can thus be derived as

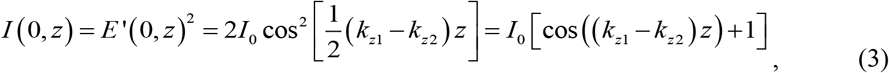

where *I*_0_ = 2*E*_1_^2^ = 2*E*_2_^2^, denoting a constant intensity distribution along the axial direction. In real applications, *I*_0_(*z*) is used because of the finite beam energy. Eq. (3) indicates that the Bessel droplet periodically repeats the intensity maxima along *z*-axis with a space interval of Δ*z* = 2π/Δ*k*_*z*_ [Fig. 1(b)], owing to the intensity modulation of the cosine function, where Δ*k*_*z*_ = *k*_*z*1_ – *k*_*z*2_. Here, Δ*k*_*z*_ is defined as the beating frequency of the two Bessel beams. The values of *k*_*z*1_ and *k*_*z*2_ are determined by the tilted angles *θ*_1_ and *θ*_2_, respectively, as shown in Fig. 1(a), which can be expressed as *k*_*z*1_ = cos(*θ*_1_)·2π*n*/*λ* and *k*_*z*2_ = cos(*θ*_2_)·2π*n*/*λ*, where *n* is the refractive index of the medium and *λ* is the laser wavelength. Based on the Fourier transform, the Bessel droplet is transformed from two concentric rings at the back focal plane of the objective, as shown in Fig. 1(b). In the imaging system, the tilted angle *θ* of the wave vector is determined by sin(*θ*_1_) = *r*_1_·*NA*/*nR* and sin(*θ*_2_) = *r*_2_·*NA*/*nR*, where *R* and *NA* are the radius of the back aperture and numerical aperture of the objective, respectively. Here, the size of the back aperture is regarded to be equal to the back focal plane of the objective. Thus, by adjusting the radii *r*_1_ and *r*_2_ of the two rings, the beating frequency Δ*k*_*z*_ can be controlled, as shown in Fig. 1(c).

**Fig. 1.**
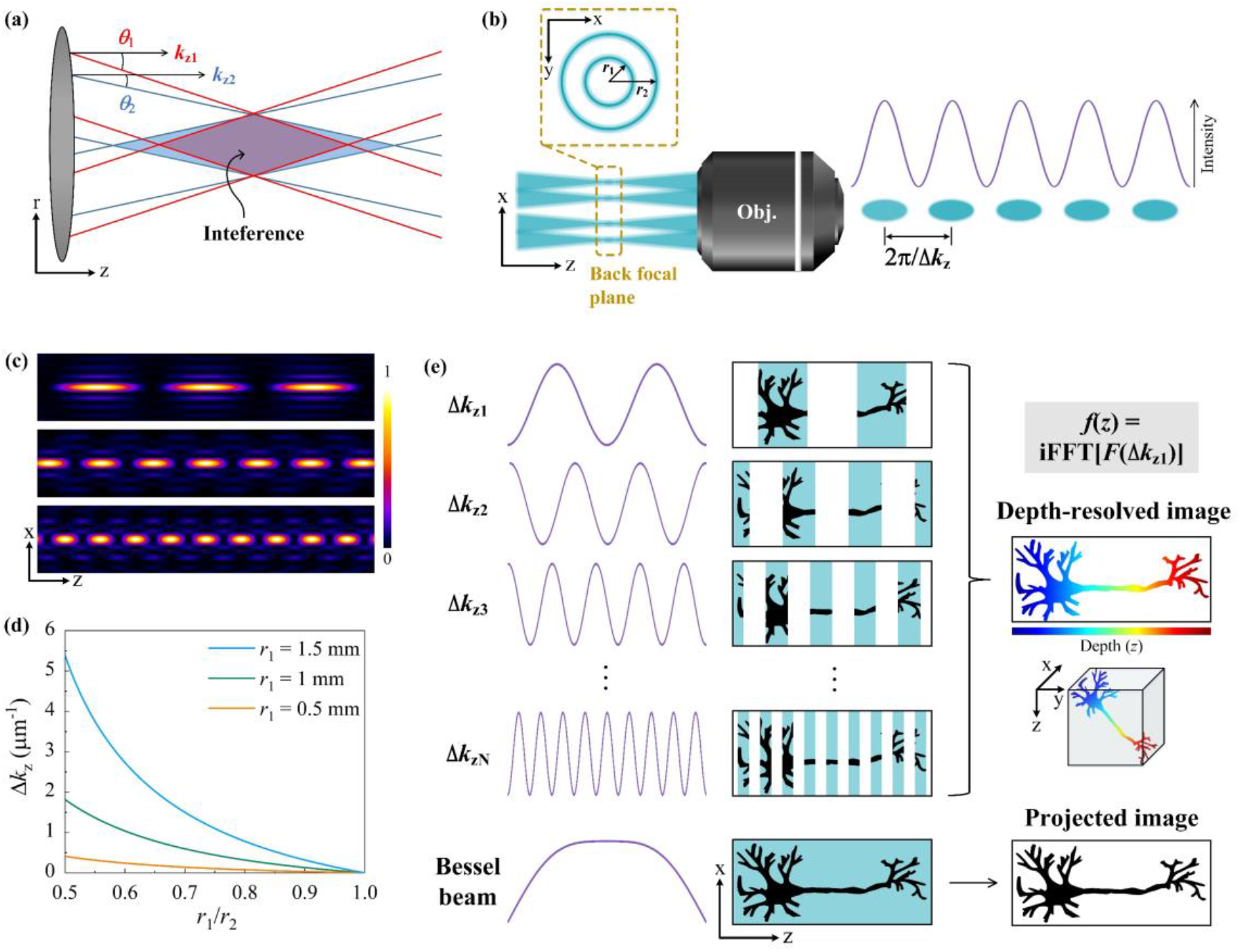
Working principle of TEMP microscopy. (a) Principle of generating the Bessel droplet. (b) Generation of the Bessel droplet in a confocal microscopy. (c) Bessel droplets with different beating frequencies. (d) Beating frequency as a function of the ratio between the radii of the inner and outer rings on the back focal plane of the objective. (e) Principle of resolving the imaging depth in the microscopy and the comparison with the conversional Bessel beam.

Fig. 1(e) demonstrates the principle to resolve the imaging depth using the illuminations of the Bessel droplets. A 3D sample is sequentially scanned by a series of Bessel droplets with different Δ*k*_*z*_. The scanning in *x*-*y* plane is controlled by galvo mirrors (GMs). As the Bessel droplets also have EDOFs as the 0th-order Bessel beam, a set of 2D projected images is obtained. Because of the modulation of cosine function on the axial intensity of the Bessel droplet, only the intensity above a certain level can excite the signal. Accordingly, evenly separated depth positions in the sample are projected to the 2D image, also following the cosine distribution. The set of cosine functions is capable to construct a Δ*k*_*z*_ space containing the depth information of the sample. By making an inverse fast Fourier transform (iFFT) from the spatial frequency domain to the spatial domain, the depth *z* can be resolved. Together with the scanning obtained *x* and *y* information, a 3D image can thus be reconstructed. In contrast, as the EDOF of the 0th-order Bessel beam has a continuous intensity distribution, the axial information is buried in its projected 2D image.

The mathematical expression of the depth-resolving process is as follows. Considering a 3D sample *f*(*x, y, z*) scanned by the Bessel droplet, a series of 2D projected images *F*(*x, y*, Δ*k*_*z*_) can be obtained by varying Δ*k*_*z*_. Thus, we can construct a spatial frequency domain containing the depth information of the sample, which is described as [27]

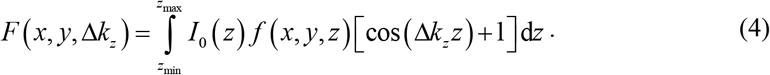

Eq. (4) shows that *z* determines the frequency of the cosine function in Δ*k*_*z*_ space, indicating that the objects at differet depths have different oscillating frequencies in Δ*k*_*z*_ space. Based on this, the axial informanton can be differentiated. The reconstructed 3D image *f*(*x, y, z*’) can be obtained through iFFT, which is contained in the following expression,

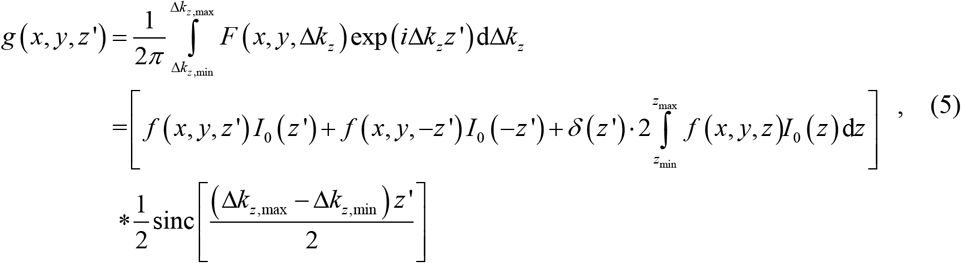

where Δ*k*_*z*_ ranges from Δ*k*_*z*,min_ to Δ*k*_*z*,max_, and “*” denotes convolution. The three terms in the square bracket on the right side of Eq. (5) represent the retrieved image, mirror image, and direct current (DC) component. The reconstructed 3D image *f*(*x, y, z*’) is achieved by normalizing to the illumination function *I*_0_(*z*’), while the mirror image and the DC component are omitted. The sinc function determines the axial resolution Δ*z*’, which is expressed in the full width at half maximum (FWHM) as

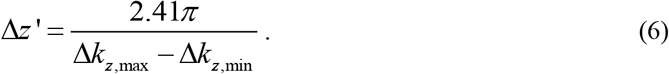

Eq. (6) indicates that a large difference between Δ*k*_*z*,max_ and Δ*k*_*z*,min_ contributes to a high axial resolution. Fig. 1(d) shows Δ*k*_*z*_ as a function of the ratio between *r*_1_ and *r*_2_. In the simulation, *R* = 4 mm, *NA* = 1.2, *n* = 1 (assuming in the air) and *λ* = 532 nm. *r*_1_ is set to be 1.5, 1, and 0.5 mm, while *r*_2_ increases from the value of *r*_1_ to its double. As shown in Fig. 1(d), a larger radius of the inner ring benefits to achieving a wider range of Δ*k*_*z*_.

### Nonlinear excitation of Bessel droplet

Except for the tunability of the beating frequency, the performance of TEMP microscopy is also largely determined by the shape of the Bessel droplet, including the “Droplet” and “Gap” regions [Fig. 2(a)]. Firstly, the lateral shape of the Droplet determines the PSF in a laser scanning confocal microscopy; Secondly, the intensity in the Gap region should be as low as possible, aiming to maximize the axial intensity gradient from the Droplet to Gap. This is critical to improve the accuracy of the iFFT process and minimize the background of the reconstructed 3D image.

**Fig. 2.**
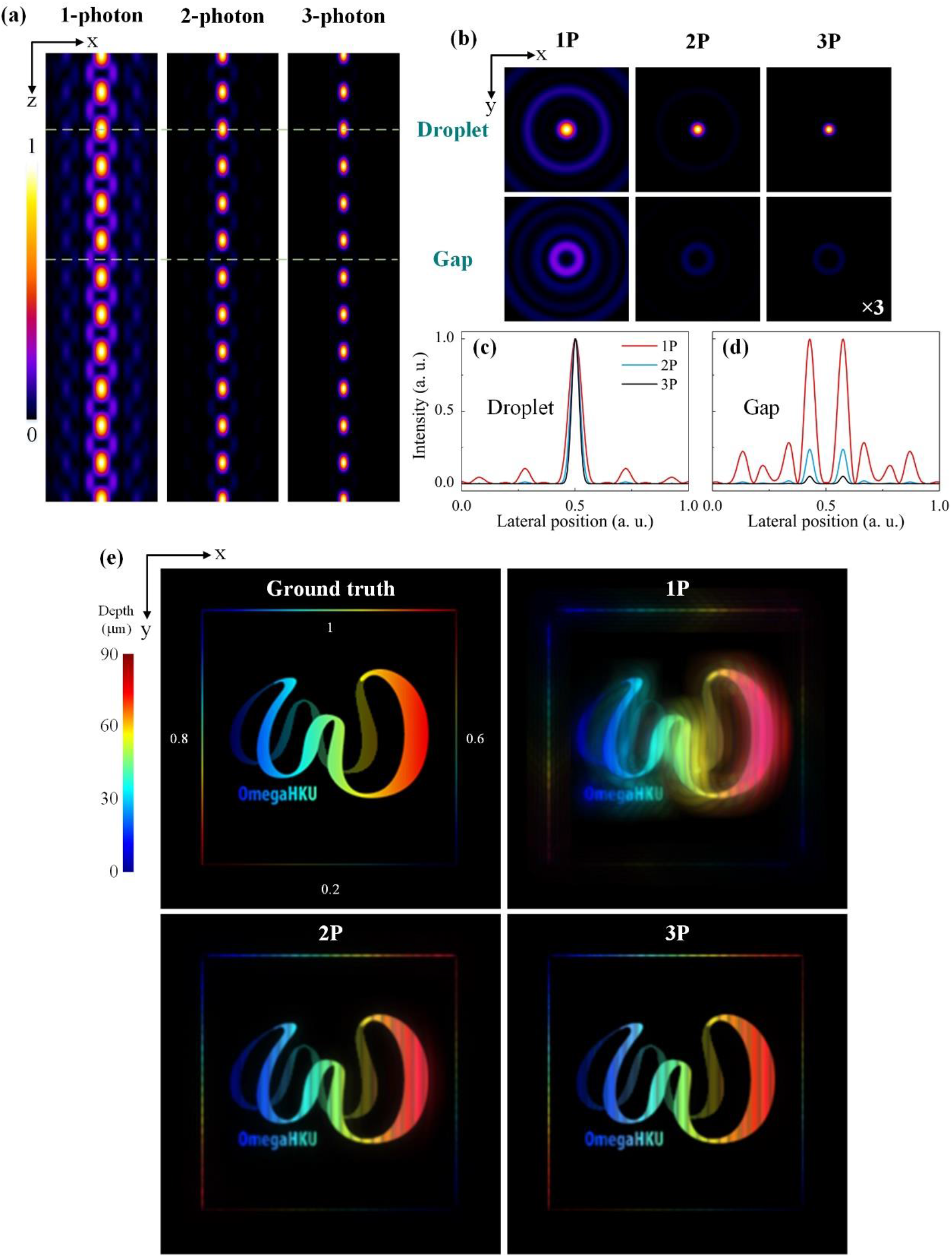
Nonlinear excitation of Bessel droplet. (a) Fluorescence intensity distributions in the *x-z* plane under the 1-, 2-, and 3-photon (1P, 2P, and 3P) excitations. (b) Lateral intensity distributions of the Droplet and Gap region. (c)(d) Intensity comparison of the Droplet and Gap. (e) Reconstruction simulation of a 3D structure under 1P, 2P, and 3P excitations.

Multiphoton excitation is an excellent approach to optimize the shape of the Bessel droplet. In contrast to the single-photon (1P) excitation, multiphoton condition is capable to confine the effective focal spot in a much smaller region owing to the higher-order nonlinearity. Assuming the dye concentration, absorption cross-section, and temporal intensity distributions of the excitation light are constants, the *m*-photon excited fluorescence can thus be simplified to [37]

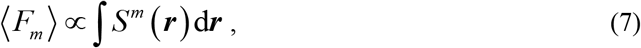

where *S*(*r*) is the spatial intensity distribution of the excitation light. Here, *S*(*r*) follows the intensity distribution of the Bessel droplet. Fig. 2(a) shows the fluorescence intensity distribution along the EDOF under the 1-, 2-, and 3-photon excitations. In the 1P case, there are clear side-lobes in the Gap region, and they periodically repeat along *z*-axis. As a comparison, the 2P and 3P cases are much cleaner in the same Gap region. Furthermore, the 3D size of the Droplet decreases with the nonlinearity, which is confirmed by Fig. 2(c). Fig. 2(b) demonstrates the lateral beam shapes of the Droplet and Gap under the three excitation situations. For the 1P excitation, there is a ring around the main spot, which will also degrade the imaging contrast as its non-ignorable intensity; multiple concentric rings exist in the Gap region of the 1P case, but they reduce to one ring with negligible intensities under the 2P and 3P conditions. As shown in Fig. 2(d), the peak intensity of the inner ring under the 2P excitation is only 24% of that under the 1P excitation, and it greatly decreases to 5% under the 3P excitation.

To demonstrate the 3D reconstruction performance under 1P, 2P, and 3P excitations, we constructed a 3D model. Fig. 2(e) shows the ground truth. The depth is coded by colors, ranging from 0 to 90 µm. The intensities of the structure vary at different axial positions, aiming to imitate the diverse intensity distributions of biological tissues. For better illustrations, a square was added around the structure, and the four sides are in different intensities (normalized intensity, 1, 0.8, 0.6, and 0.2). The value is labeled near each side. For the scanning of the Bessel droplets, the beating frequency ranges from 0 to 1 µm^-1^. Comparing the three depth-resolved images, the structure is extremely burred under the 1P excitation, mainly resulting from the side-lobes in the Gap and Droplet regions of the Bessel droplet. In contrast, the 2P and 3P cases are much clearer, where the twin “*ω*” shapes are well reconstructed approaching to the ground truth. More importantly, the multi-photon excitation also demonstrates a better reconstruction of the line with weak intensities, indicating its powerful ability to resolve the weak fluorescence structures in the bio-tissues, like the thin neurons in the mouse brain.

## Imaging performance

### Bessel droplet PSF in 2P microscopy

According to the simulation results [Fig. 2(e)], as the imaging performances are similar between the 2P and 3P cases, we experimentally demonstrate the TEMP microscopy under 2P excitation in the following sections. To demonstrate the tunability of the Bessel droplets in the TEMP microscopy, the PSFs of different beating frequencies were measured. The axial beam shape was scanned by fluorescence microspheres (1 µm, F8819, Life Technologies Ltd.). The axial scan steps for the 0th-order Bessel beam and Bessel droplets were 2 and 1 µm, respectively. Figs. 3(a) and 3(b) demonstrate the beam shapes in *x-z* plane and the corresponding axial intensity distributions, respectively, when the beating frequency is set to be 0 (0th-order Bessel beam), 0.25, 0.5, and 0.75 µm^-1^. As shown in Fig. 3(a), the axial repetition of the Droplet pattern varies with the beating frequency. Furthermore, the 2P-excited Bessel droplets have few side lobes, both for Droplet and Gap regions, which agrees well with the simulations [Fig. 2]. The DOF of the 0th-order Bessel beam is 54 µm [Fig. 3(b)]. As the optical beating happens within the DOFs of both Bessel beams, the peak intensities of the Droplet spots in each Bessel droplet are under a Gaussian-like envelope for Δ*k*_*z*_ > 0. The spatial intervals of the Droplet spots were measured to be 30, 15, and 10 µm corresponding to the beating frequency of 0.25, 0.5, and 0.75 µm^-1^, respectively. Their respective products (Δ*z* × Δ*k*_*z*_) are all equal to 7.5, indicating a precise periodic modulation of the Bessel droplets. The product in experiments is slightly larger than the theoretical value 2*π* (assuming in the air), resulting from the refractive index of the medium between the objective and sample.

**Fig. 3.**
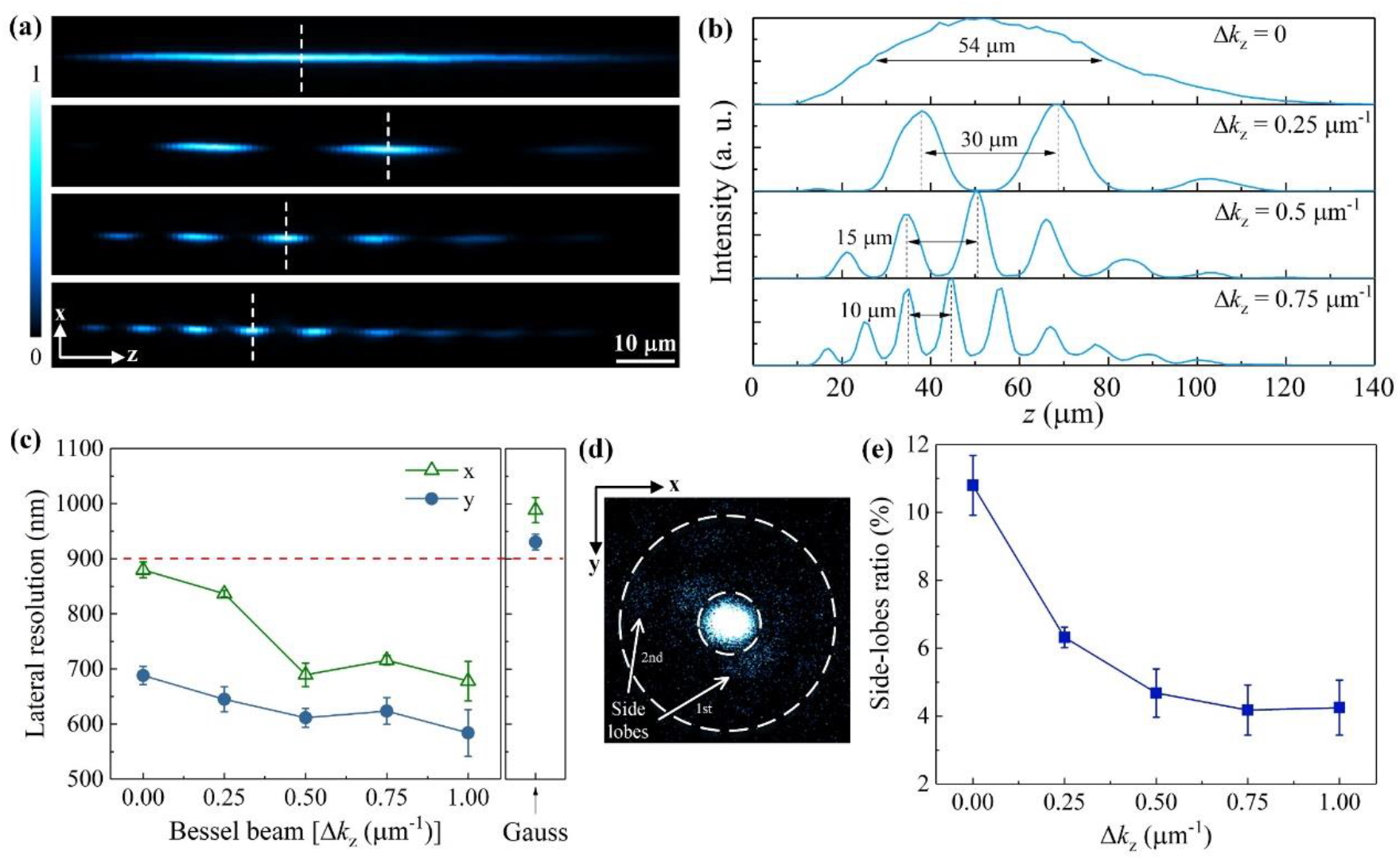
Bessel droplet PSF in 2P microscopy. (a) Axial beam shapes and (b) intensity distributions of the 0th-order Bessel beam (Δ*k*_*z*_ = 0) and Bessel droplet at Δ*k*_*z*_ = 0.25, 0.5, and 0.75 µm^-1^. (c) Lateral resolution of Bessel droplet as a function of the beating frequency and the lateral resolution of Gaussian beam. (d) Lateral shape of the 0th-order Bessel beam. (e) Intensity ratio between the side lobes and main spot as a function of the beating frequency.

The lateral resolutions are compared among the Bessel droplet, Gaussian beam, and 0th-order Bessel beam (Δ*k*_*z*_ = 0). The beams were laterally scanned by 200-nm fluorescence microspheres (F8809, Life Technologies Ltd.). Their corresponding axial positions are labeled by the white dashed lines in Fig. 3(a). As shown in Fig. 3(c), the PSF tends to decrease when increasing the beating frequency. The lateral beam sizes of the 0th-order Bessel beam are 880 ± 15 nm (*x*-axis) and 688 ± 17 nm (*y*-axis), while those of the Bessel droplet are 678 ± 36 nm (*x*-axis) and 584 ± 42 nm (*y*-axis) when Δ*k*_*z*_ = 1 µm^-1^. The improvements are around 23% (*x*-axis) and 15% (*y*-axis). At the same focusing condition, the lateral resolutions of the Gaussian beam are 989 ± 23 nm (*x*-axis) and 930 ± 14 nm (*y*-axis). The resolutions of the 0th-order Bessel beam and Bessel droplet are both better than the Gaussian beam, resulting from their ring-shaped beams at the back focal plane of the objective. Only the high spatial frequency components are focused, making the lateral beam size smaller at the focal region of the objective. On the other hand, owing to the interference of two Bessel beams, the side lobes of the Bessel droplet can be diminished. Fig. 3(d) shows the lateral beam shape of the 0th-order Bessel beam, where the 1st and 2nd side-lobes are around the main spot. By integrating the intensities of the side lobes and main spot, respectively, we calculated the intensity ratio between the two regions as a function of the beating frequency, as shown in Fig. 3(e). The side-lobes ratio decreases when increasing the Δ*k*_*z*_. The ratio is ∼11% for the 0th-order Bessel beam, while it reduces to ∼4% when Δ*k*_*z*_ = 1 µm^-1^.

The smaller lateral beam sizes of the Bessel droplets will make the lateral image resolution higher than that obtained from the conventional Bessel and Gaussian beams. Moreover, the side-lobes suppression of the Bessel droplets contributes to a cleaner image background, which will enhance the imaging contrast. Together with the depth-resolving capacity, the Bessel droplets have distinct advantages over the 0th-order Bessel and Gaussian beams, thus can achieve high-resolution and high-contrast images.

### Highly scattering microspheres phantom

A primary advantage of MPM is its unique ability to penetrate highly scattering tissues because of its nonlinear light–matter interaction. On the other hand, the diffraction-free and self-healing nature of the Bessel beams also features strong penetration ability compared with the conventional Gaussian beam. As the Bessel droplet is generated by the interference of two 0th-order Bessel beams, it owns the nature of the Bessel beam. The TEMP microscopy integrates the multiphoton excitation and NDB techniques, thus it has great potential to enhance the penetration capacity. To test the penetration performance of the TEMP microscopy, we prepared a highly scattering microspheres phantom. The sample preparation is described in the experimental section. The phantom was sequentially scanned by the Bessel droplets with Δ*k*_*z*_ ranging from 0 to 1 µm^-1^. The depth-resolved image from the Bessel-droplet stacks is shown in Fig. 4(a), where the color code represents the axial position. The axial resolution of the reconstructed Bessel droplet image was calculated to be ∼3.8 µm based on Eq. (6). The same region of the sample was then scanned by the Gaussian beam layer by layer with an axial step of 3 µm (the DOF of the Gaussian beam is 4 µm, as shown in Fig. S2), where an electrical actuator was used to control the axial shift of the sample. Fig. 4(b) shows the color-coded Gaussian stacks in a projected image. The color bar is the same for both Figs. 4(a) and 4(b). Comparing the color distributions of the two images, the depth-resolved image in TEMP microscopy has an excellent consistency with the standard Gaussian results. The image obtained from the 0th-order Bessel beam is shown in Fig. 4(c), where a strong background exists, especially around the beads. This is attributed to the scattering nature of the phantom and the side lobes of the Bessel beam.

**Fig. 4.**
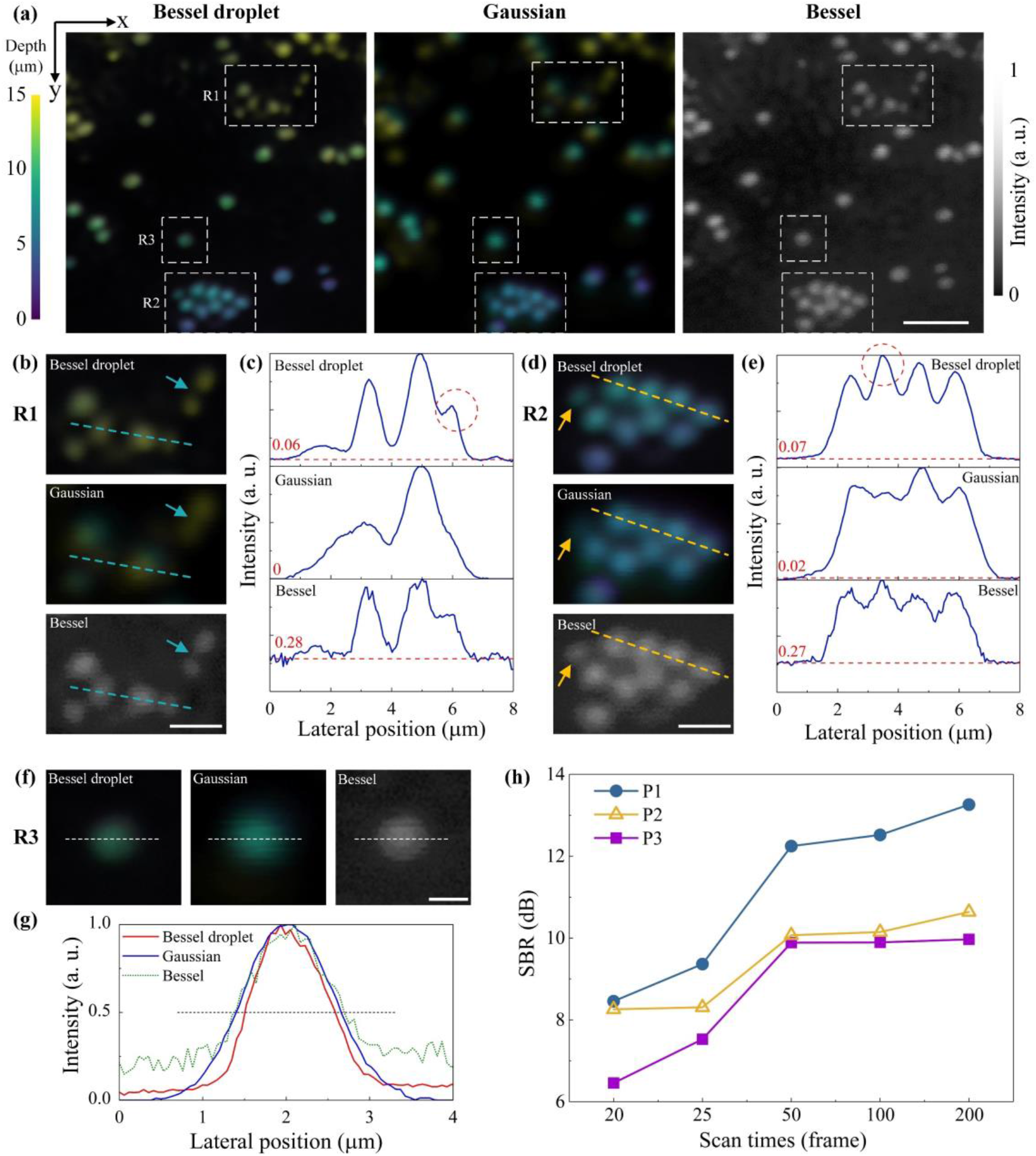
Highly scattering microspheres phantom. (a) Depth-resolved image from the Bessel droplet stacks, projected image of the Gaussian stacks, and the image obtained by the 0th-order Bessel beam for a highly scattering microspheres phantom. (b-g) Comparisons of the imaging performance at region (b)(c) R1, (d)(e) R2, and (f)(g) R3. Scale bar: (a) 5 µm, (b)(d) 2 µm, and (f) 1 µm. (h) SBR of the reconstructed image under different scan times of the Bessel droplets. Post-obj. power: ∼80 mW (Bessel droplet and Bessel beam), ∼16 mW (Gaussian beam). Visualization 1 demonstrates the image stacks comparison between the Bessel droplet and Gaussian beam.

Except for the depth-resolving capacity, we also compared the resolution and contrast of the three images. Three regions (R1, R2, and R3) located at different depths in Fig. 4(a) are magnified for the comparisons. As shown in Fig. 4(b), more beads are resolved in the Bessel droplet image than that in the Gaussian image, meanwhile the beads in the Bessel droplet image demonstrate much clearer shapes. There are seven microspheres in total presented in Bessel droplet and Bessel images, but only around five microspheres are shown in the Gaussian image. As indicated by the arrows, the two beads are separated well in the Bessel droplet and Bessel images, but they cannot be resolved by the Gaussian beam. A quantitative intensity comparison of the dashed line is shown in Fig. 4(c). Three peaks can be clearly observed in the Bessel droplet image, and the third peak is not resolved well by the Bessel beam. However, only two peaks are shown in the Gaussian image. The red baseline illustrates the background intensity in each image [Fig. 4(c)]. The Bessel image has the most serious background (0.28, normalized to the maximum intensity), while the Bessel droplet (0.06) and Gaussian (∼0) images are much lower. The strong image background from the 0th-order Bessel beam is attributed to its intense side lobes. In contrast, the side-lobes intensities of different Bessel droplets are all lower than those of the 0th-order Bessel beam [Fig. 3(e)], thus the reconstructed image has a cleaner background, and its contrast performance approaches to the standard Gaussian image.

The same performance can be confirmed in region R2. As shown in Fig. 4(d), a microsphere can hardly be seen in the Gaussian image (pointed by the arrow), but it is resolved well in the other two images, indicating the excellent non-diffracting nature of the Bessel beam group. As shown in Fig. 4(e), the 2nd peak in the Gaussian image is hardly resolved. For the background intensities of the three images, they are consistent with region R1. Figs. 4(f) and 4(g) demonstrate the comparison of a single microsphere, corresponding to region R3. The FWHM of the microsphere in the Bessel droplet image is the smallest, while the FWHMs from the other two images are similar. This is ascribed to the smallest PSF of the Bessel droplet among the three beams. In contrast to the Bessel and Gaussian beam, the TEMP microscopy can achieve a better resolution and contrast in the highly scattering samples, more importantly, with the excellent depth-resolving ability.

We also investigated the depth-resolving performance by varying the scan times by the Bessel droplets. By fixing the range of Δ*k*_*z*_ (0 ∼ 1 µm^-1^), we obtained the reconstructed images under 20, 25, 50, 100, and 200 raw images. The 3D reconstruction from 200 raw images is shown in Fig. 4(a), and the other reconstructed images are shown in Fig. S3. The SBRs of these reconstructed images are compared by measuring the SBRs of the beads located at different depths. Fig. 4(h) shows the image SBR as a function of the scan times by the Bessel droplets. According to the three positions (P1, P2, and P3, shown in Fig. S3), the image SBR increases when using more raw images. This is attributed to the gained exposure time from more scans. Notably, the reconstructed image from 20 raw data still has a SBR above 6.5 dB, which is also beyond acceptable for applications. Moreover, we demonstrated the depth-resolving performance at different Δ*k*_*z*_ ranges. By fixing the number of raw images (50 frames), we compared the reconstructed images at Δ*k*_*z*_ from 0 to 1 and 0 to 0.25 µm^-1^ [Fig. S3]. When the range of Δ*k*_*z*_ increases, the axial resolution is also improved, which is consistent with Eq. (6).

The axial scanning speed is compared between the TEMP microscopy and the conventional Gaussian scanning. The SLM refresh rate determines the axial scan rate of the Bessel droplet, which is 60 Hz in this experiment; while the sample is axially shifted by the electrical actuator in the Gaussian module, and its speed is up to ∼10 Hz. Considering the raw images obtained for the Bessel droplet (20 frames) and Gaussian beam (6 frames) to achieve a similar axial resolution and image SBR, the axial scanning speed of the TEMP microscopy is 1.8 times of that with the Gaussian beam. Notably, if applying a high-speed DMD, the speed can be further boosted.

### Mouse brain neurons

To demonstrate the applicability of TEMP microscopy for biological tissues, a 50-µm-thick mouse brain tissue was prepared for imaging. The imaging procedure was the same as the microspheres phantom. The depth-resolved image from the Bessel droplets, Gaussian projected image, and image obtained by the 0th-order Bessel beam are demonstrated in Fig. 5(a). The neurons extend within the thickness of 26 µm. Comparing the Bessel droplet and Gaussian images, the axial positions of the neurons are well matched, indicated by the colors. For example, the neurons in blue and red locate at the shallow and deep positions, respectively. They are well resolved along the axial direction by the Bessel droplets with an axial resolution approaching to the Gaussian beam. Comparing the three images, some thin neurons with weak fluorescence in the Gaussian image can hardly be seen, while they are demonstrated in the other two images. Figs. 5(b) and 5(c) show the intensity comparison of the neurons in two regions. In Fig. 5(b), the four peaks are distinguishable in the Bessel droplet image, while Peak 2 and Peak 3 are missing in the Gaussian image. As the background is serious in the Bessel image, the four peaks are all buried in noises. The same performance is also shown in Fig. 5(c), Peak 2 can only be observed in the Bessel droplet image. The magnified images of the corresponding region are demonstrated in Fig. 5(d). As pointed by the arrows, the weak fluorescence neurons are resolved by the Bessel droplets, while they cannot be seen in the Gaussian image and are blurred in the Bessel image.

**Fig. 5.**
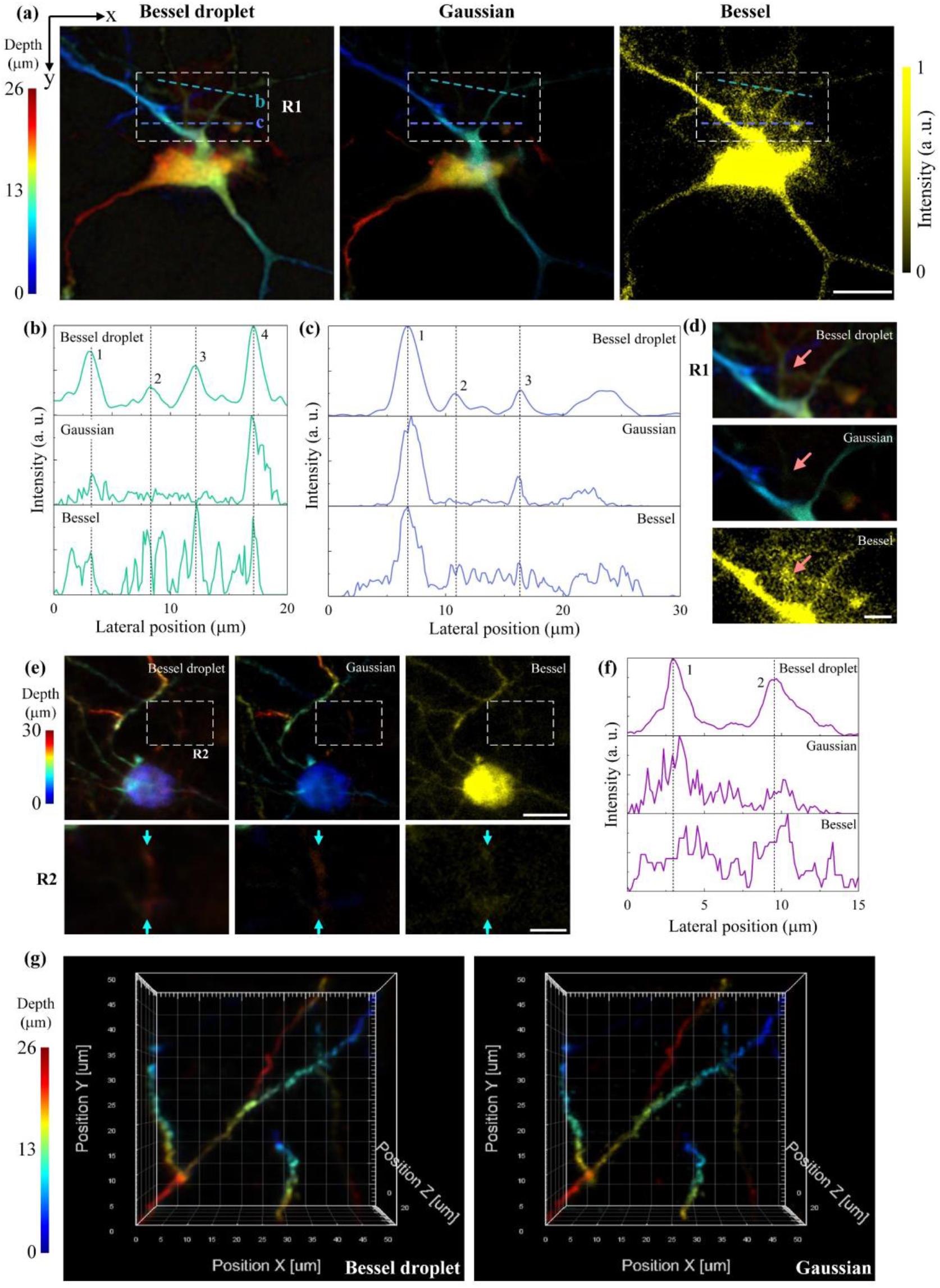
Mouse brain neurons. (a) Depth-resolved image from the Bessel droplet stacks, projected image of the Gaussian stacks, and the image obtained by the 0th-order Bessel beam for a mouse brain slice. (b-d) Comparisons of the imaging performance at region R1. (e) Images obtained in the three modalities and their corresponding magnified images of region R2. (f) Intensity comparison of the same region. (g) 3D demonstration of the dendrites in the mouse brain. Scale bar: (a)(e, top row) 15 µm and (d)(e, bottom row) 5 µm. Post-obj. power: ∼70 mW (Bessel droplet and Bessel beam), ∼13 mW (Gaussian beam). Visualization 2 and 3 demonstrate the image stacks comparison between the Bessel droplet and Gaussian beam for the neurons in (g) and (e), respectively.

The excellent resolving ability of the Bessel droplet is attributed to its better lateral resolution, less side-lobes, and more importantly, its diffraction-free and self-healing nature. Owing to this unique nature, the Bessel droplet and 0th-order Bessel beam can both image these weak fluorescence neurons, while the Gaussian beam cannot. Because of the first two advantages, the Bessel droplets have a better discrimination capacity over the 0th-order Bessel beam. Multiple illuminations on the same axial position of the sample using the Bessel droplets are another advantage over the Gaussian beam. As multiple illuminations increase the exposure time of the images while maintaining low illumination levels, it lowers the risks of photodamaging and photobleaching on samples, as well as enhancing the image SBR, avoiding directly increasing the excitation intensity like the Gaussian beam.

The outstanding performance of the TEMP microscopy is confirmed in other regions of the brain slice. As shown in Fig. 5(e), not only the depth positions of the neurons are well resolved by the Bessel droplets, but its magnified image of region R2 also demonstrates more neuron details. The intensity distributions of this region are shown in Fig. 5(f). The two peaks are clearly revealed in the Bessel droplet image, compared with the other two modalities. Fig. 5(g) shows a 3D demonstration of a brain region with dendrites. The depth-resolved image has an excellent consistency with the standard result obtained from the Gaussian beam.

## Discussions

Here, we demonstrated a novel and powerful tomographic-encoded MPM using the Bessel droplets, termed TEMP microscopy. It allowed fast axial scanning, less risks of photodamage and photobleaching, and high-resolution and high-contrast imaging in MPM.

We realized the axial scanning in MPM by sweeping the Bessel droplets with different beating frequencies. This method is inertia-free during axial scanning, avoiding the slow mechanical shift (up to ∼10 Hz) of the sample or objective. The axial scanning speed was mainly dependent on the refresh rate of the SLM (typically 60 Hz). The speed had a great potential to be boosted by replacing a high-speed DMD (up to 20 kHz). Expect for the speed, this approach also got rid of the focus aberrations, which was usually induced by the ETL, TAG lens, and reverberation microscopy [15], as they introduced only a quadratic phase function for re-focusing. In contrast, the TEMP microscopy took advantages of the non-diffracting and self-healing nature of the Bessel beam, which provided us a uniform lateral resolution along the depth, as well as the scattering-against property.

The multiphoton excitation empowered the Bessel droplet with higher “Droplet” purity (Fig. S4), which greatly improved the depth-resolving accuracy and the quality of the reconstructed 3D image. Given that the quantity of the raw images obtained from the Bessel droplets determined the SBR of the reconstructed image and the image SBR under multiphoton excitations was inherently better than that under 1-photon excitation, fewer raw images (∼20 images) were required for the image reconstruction to achieve an acceptable SBR in the TEMP microscopy.

Compared with the conventional 0th-order Bessel beam, the side lobes of the Bessel droplet had been efficiently suppressed. This is beneficial to enhance the image contrast, as the side-lobes-induced background was minimized. Notably, less average power from the side lobes was illuminated on samples, alleviating the risks of photodamage and photobleaching in the MPM. In contrast to the Gaussian beam, the Bessel droplet utilized an alternative way to improve the image SBR, that is, by increasing the exposure time for each axial position while maintaining low illumination levels, avoiding directly increasing the illumination power on fluorescence samples that may raise the risks of photodamage and photobleaching. The gain of the exposure time from the Bessel droplets was an inherent merit of the TEMP technique, as multiple axial positions were simultaneously illuminated by each Bessel droplet. On the contrary, the Gaussian beam focused on only one layer at each scanning.

Apart from the image contrast, the later resolution of the Bessel droplet is also better than the 0th-order Bessel beam and Gaussian beam, and its axial resolution approached to the Gaussian beam under the same focusing conditions. According to the imaging performance in highly scattering microspheres phantom and mouse brain slice, the resolved axial positions in the TEMP microscopy had excellent consistencies with those obtained by the Gaussian beam. More importantly, owing to the excellent resolution, contrast, and non-diffracting properties of the Bessel droplet, more sample details can be visualized by the Bessel droplet in contrast to the other two modalities.

The TEMP technique is also an easy-plug-in approach for current confocal scanning microscopy. Compared with other axial scan technologies reported in recent years, which required complex beam alignments [16], laser pulses multiplexing [15], multiple laser paths for different layer excitations [17, 38], multiple detectors [18], or special designed optical structures [19], the TEMP microscopy only required a SLM or DMD to insert as a reflecting mirror in the beam path of the excitation laser, which saved much efforts. We believe that the TEMP microscopy holds great potential for fast volumetric multiphoton imaging, especially for highly scattering tissues.

## Materials and methods

### Experimental setup

Fig. S1 shows the experimental setup for the TEMP microscopy. The excitation light was from a customized mode-locked fiber laser (1065 nm, 30 MHz, and 180 fs) for 2P excitation. The system can be switched between the Bessel route (flip on M1 and M3) and Gaussian route. The Bessel droplet and Bessel beam were generated by the SLM (Holoeye, PLUTO). Visualization 4 demonstrated the variation of the Bessel droplets controlled by different phase masks. The focal lengths of the lens are *f*_1_ = 200 mm, *f*_2_ = 250 mm, *f*_3_ = 125 mm, *f*_4_ = 80 mm, and *f*_5_ = 150 mm. A water-immersion objective (60×, 1.2 NA, UPlanSpo, Olympus) was used to focus the excitation light into the sample. The diameter of the back aperture of the objective was ∼8 mm. The diameter of the collimated Gaussian beam on the back aperture of the objective was 4 mm, thus the effective NA for the Gaussian beam was ∼0.6. The diameter of the inner ring of the Bessel droplet was fixed to ∼3 mm, while the outer ring was increased from ∼3 to ∼4.4 mm. Thus, the maximum effective NA for the Bessel droplet was ∼0.66. Therefore, the two beams were in similar focus conditions. The *x*-*y* laser scanning was achieved by a pair of GMs (Cambridge Technology 6220H). The excited fluorescence signal was collected in the forward direction by a condenser lens (Olympus U-AAC, NA 1.4) and detected by a PMT (Hamamatsu, H10723-20). The residual excitation beam was cleaned up by two shortpass filters (Semrock, BSP01-785R-25) before the PMT. The electrical signal from the PMT was subsequently digitized by an analog-to-digital (A/D) converter card (National Instruments PCI-6110, 5 MS/sec). The control of the entire imaging system, including laser scanning, data collection, and image reconstruction, was realized by a MATLAB-based multifunctional program. It should be pointed out that the current MPM operated in transmission mode with forward detection, while it could easily be modified to implement epi-detection.

### Sample preparation

Highly scattering microspheres phantom: fluorescence beads (1 µm, F8819, Life Technologies Ltd.) and non-absorptive titanium dioxide (TiO_2_) nanospheres (diameter 10 nm) were embedded in a cured agarose gel phantom. The TiO_2_ nanospheres performed as an optical scattering medium akin to most biological tissues. The standard fluorescence beads were adopted to assure uniform signal intensity at different depths of the phantom.

Mouse brain neurons: The mouse expresses the yellow fluorescent protein (YFP) in a subset of motor and sensory neurons (Thy1-YFP H-line). Layer-V pyramidal neurons in prefrontal cortex were imaged in this paper. All experiments with these samples were approved and performed in accordance with institutional guidelines of the University of Hong Kong.

## Author Contributions

†These authors contributed equally to this work (H.H. and X.D.). H.H. and X.D. conceived the idea, performed the research, and analyzed the data. C.S.W.L provided the biological sample. Y.R., C.S.W.L, K.K.T, and K.K.Y.W. supervised the whole project. H.H. wrote the manuscript with input from all authors. All authors discussed and revised the manuscript.

## Data availability

All the data supporting our findings are presented in the main text and the supplemental document. The MATLAB codes are available from the corresponding author upon reasonable request.

## References

1. J. Wu, N. Ji, and K. K. Tsia, “Speed scaling in multiphoton fluorescence microscopy,” Nat. Photonics 15, 800–812 (2021).

2. N. G. Horton, K. Wang, D. Kobat, C. G. Clark, F. W. Wise, C. B. Schaffer, and C. Xu, “In vivo three-photon microscopy of subcortical structures within an intact mouse brain,” Nat. Photonics 7, 205–209 (2013).

3. T. Wang and C. Xu, “Three-photon neuronal imaging in deep mouse brain,” Optica 7, 947–960 (2020).

4. Y. Hontani, F. Xia, and C. Xu, “Multicolor three-photon fluorescence imaging with single-wavelength excitation deep in mouse brain,” Sci. Adv. 7, eabf3531 (2021).

5. J. Wu, Y. Liang, S. Chen, C.-L. Hsu, M. Chavarha, S. W. Evans, D. Shi, M. Z. Lin, K. K. Tsia, and N. Ji, “Kilohertz two-photon fluorescence microscopy imaging of neural activity in vivo,” Nat. Methods 17, 287–290 (2020).

6. K. Choe, Y. Hontani, T. Wang, E. Hebert, D. G. Ouzounov, K. Lai, A. Singh, W. Béguelin, A. M. Melnick, and C. Xu, “Intravital three-photon microscopy allows visualization over the entire depth of mouse lymph nodes,” Nat. Immunol. 23, 330–340 (2022).

7. B. Sun, M. Wang, A. Hoerder-Suabedissen, C. Xu, A. M. Packer, and F. G. Szele, “Intravital Imaging of the Murine Subventricular Zone with Three Photon Microscopy,” Cereb. Cortex (2022).

8. F. Helmchen and W. Denk, “Deep tissue two-photon microscopy,” Nat. Methods 2, 932–940 (2005).

9. J. N. Stirman, I. T. Smith, M. W. Kudenov, and S. L. Smith, “Wide field-of-view, multi-region, two-photon imaging of neuronal activity in the mammalian brain,” Nat. Biotechnol. 34, 857–862 (2016).

10. T. Ragan, L. R. Kadiri, K. U. Venkataraju, K. Bahlmann, J. Sutin, J. Taranda, I. Arganda-Carreras, Y. Kim, H. S. Seung, and P. Osten, “Serial two-photon tomography for automated ex vivo mouse brain imaging,” Nat. Methods 9, 255–258 (2012).

11. C. Li, J. Shi, X. Wang, B. Wang, X. Gong, L. Song, and K. K. Wong, “High-energy all-fiber gain-switched thulium-doped fiber laser for volumetric photoacoustic imaging of lipids,” Photonics Res. 8, 160–164 (2020).

12. B. Berge and J. Peseux, “Variable focal lens controlled by an external voltage: An application of electrowetting,” Eur. Phys. J. E 3, 159–163 (2000).

13. A. Mermillod-Blondin, E. McLeod, and C. B. Arnold, “High-speed varifocal imaging with a tunable acoustic gradient index of refraction lens,” Opt. Lett. 33, 2146–2148 (2008).

14. M. Duocastella, B. Sun, and C. B. Arnold, “Simultaneous imaging of multiple focal planes for three-dimensional microscopy using ultra-high-speed adaptive optics,” J. Biomed. Opt. 17, 050505 (2012).

15. D. R. Beaulieu, I. G. Davison, K. Kılıç, T. G. Bifano, and J. Mertz, “Simultaneous multiplane imaging with reverberation two-photon microscopy,” Nat. Methods 17, 283–286 (2020).

16. J. Demas, J. Manley, F. Tejera, K. Barber, H. Kim, F. M. Traub, B. Chen, and A. Vaziri, “High-speed, cortex-wide volumetric recording of neuroactivity at cellular resolution using light beads microscopy,” Nat. Methods 18, 1103–1111 (2021).

17. S. Weisenburger, F. Tejera, J. Demas, B. Chen, J. Manley, F. T. Sparks, F. M. Traub, T. Daigle, H. Zeng, and A. Losonczy, “Volumetric Ca2+ imaging in the mouse brain using hybrid multiplexed sculpted light microscopy,” Cell 177, 1050-1066. e1014 (2019).

18. A. Badon, S. Bensussen, H. J. Gritton, M. R. Awal, C. V. Gabel, X. Han, and J. Mertz, “Video-rate large-scale imaging with Multi-Z confocal microscopy,” Optica 6, 389–395 (2019).

19. T. Chakraborty, B. Chen, S. Daetwyler, B.-J. Chang, O. Vanderpoorten, E. Sapoznik, C. F. Kaminski, T. P. Knowles, K. M. Dean, and R. Fiolka, “Converting lateral scanning into axial focusing to speed up three-dimensional microscopy,” Light-Sci. Appl. 9, 1–12 (2020).

20. R. Lu, W. Sun, Y. Liang, A. Kerlin, J. Bierfeld, J. D. Seelig, D. E. Wilson, B. Scholl, B. Mohar, and M. Tanimoto, “Video-rate volumetric functional imaging of the brain at synaptic resolution,” Nat. Neurosci. 20, 620–628 (2017).

21. R. Lu, Y. Liang, G. Meng, P. Zhou, K. Svoboda, L. Paninski, and N. Ji, “Rapid mesoscale volumetric imaging of neural activity with synaptic resolution,” Nat. Methods 17, 291–294 (2020).

22. X.-J. Tan, C. Kong, Y.-X. Ren, C. S. Lai, K. K. Tsia, and K. K. Wong, “Volumetric two-photon microscopy with a non-diffracting Airy beam,” Opt. Lett. 44, 391–394 (2019).

23. H. He, C. Kong, X.-J. Tan, K. Y. Chan, Y.-X. Ren, K. K. Tsia, and K. K. Wong, “Depth-resolved volumetric two-photon microscopy based on dual Airy beam scanning,” Opt. Lett. 44, 5238–5241 (2019).

24. G. Thériault, M. Cottet, A. Castonguay, N. McCarthy, and Y. De Koninck, “Extended two-photon microscopy in live samples with Bessel beams: steadier focus, faster volume scans, and simpler stereoscopic imaging,” Front. Cell. Neurosci. 8, 139 (2014).

25. H. He, Y.-X. Ren, R. K. Chan, W. So, H. K. Fok, C. S. Lai, K. K. Tsia, and K. K. Wong, “Background-free volumetric two-photon microscopy by side-lobes-cancelled bessel beam,” IEEE J. Sel. Top. Quantum Electron. 27, 1–7 (2021).

26. H. He, C. Kong, K. Y. Chan, W. So, H. K. Fok, Y.-X. Ren, C. S. Lai, K. K. Tsia, and K. K. Wong, “Resolution enhancement in an extended depth of field for volumetric two-photon microscopy,” Opt. Lett. 45, 3054–3057 (2020).

27. L. Gong, S. Lin, and Z. Huang, “Stimulated Raman Scattering Tomography Enables Label-Free Volumetric Deep Tissue Imaging,” Laser Photon. Rev. 15, 2100069 (2021).

28. A. Song, A. S. Charles, S. A. Koay, J. L. Gauthier, S. Y. Thiberge, J. W. Pillow, and D. W. Tank, “Volumetric two-photon imaging of neurons using stereoscopy (vTwINS),” Nat. Methods 14, 420–426 (2017).

29. X. Chen, C. Zhang, P. Lin, K.-C. Huang, J. Liang, J. Tian, and J.-X. Cheng, “Volumetric chemical imaging by stimulated Raman projection microscopy and tomography,” Nat. Commun. 8, 1–12 (2017).

30. J. Tang, J. Ren, and K. Y. Han, “Fluorescence imaging with tailored light,” Nanophotonics 8, 2111–2128 (2019).

31. X. Hua, C. Guo, J. Wang, D. Kim-Holzapfel, B. Schroeder, W. Liu, J. Yuan, J. French, and S. Jia, “Depth-extended, high-resolution fluorescence microscopy: whole-cell imaging with double-ring phase (DRiP) modulation,” Biomed. Opt. Express 10, 204–214 (2019).

32. W. Chaze, O. Caballina, G. Castanet, and F. Lemoine, “The saturation of the fluorescence and its consequences for laser-induced fluorescence thermometry in liquid flows,” Exp. Fluids 57, 1–18 (2016).

33. J.-Y. Tinevez, J. Dragavon, L. Baba-Aissa, P. Roux, E. Perret, A. Canivet, V. Galy, and S. Shorte, “A quantitative method for measuring phototoxicity of a live cell imaging microscope,” in Methods Enzymol. (Elsevier, 2012), pp. 291–309.

34. J. Icha, M. Weber, J. C. Waters, and C. Norden, “Phototoxicity in live fluorescence microscopy, and how to avoid it,” Bioessays 39, 1700003 (2017).

35. J. Arlt and K. Dholakia, “Generation of high-order Bessel beams by use of an axicon,” Opt. Commun. 177, 297–301 (2000).

36. D. B. Ruffner and D. G. Grier, “Optical conveyors: a class of active tractor beams,” Phys. Rev. Lett. 109, 163903 (2012).

37. C. Xu and W. W. Webb, “Multiphoton excitation of molecular fluorophores and nonlinear laser microscopy,” in Topics in fluorescence spectroscopy (Springer, 2002), pp. 471–540.

38. A. Cheng, J. T. Gonçalves, P. Golshani, K. Arisaka, and C. Portera-Cailliau, “Simultaneous two-photon calcium imaging at different depths with spatiotemporal multiplexing,” Nat. Methods 8, 139–142 (2011).

